# Metheor: Ultrafast DNA methylation heterogeneity calculation from bisulfite read alignments

**DOI:** 10.1101/2022.07.20.500893

**Authors:** Dohoon Lee, Bonil Koo, Jeewon Yang, Sun Kim

## Abstract

**Motivation:** Phased DNA methylation states within bisulfite sequencing reads are valuable source of information that can be used to estimate epigenetic diversity across cells as well as epigenomic instability in individual cells. Various measures capturing the heterogeneity of DNA methylation states have been proposed for a decade. However, in routine analyses on DNA methylation, this heterogeneity is often overlooked by computing average methylation levels at CpG sites. In this study, to facilitate the application of the DNA methylation heterogeneity measures in downstream epigenomic analyses, we present a Rust-based, extremely fast and lightweight bioinformatics toolkit called Metheor.

**Results:** We benchmark the performance of Metheor against existing code implementation for DNA methylation heterogeneity measures in three different scenarios of simulated bisulfite sequencing datasets. Metheor was shown to dramatically reduce the execution time up to 300-fold and memory footprint up to 60-fold, while producing the same results with the original implementation.

**Availability:** Source code for Metheor is at https://github.com/dohlee/metheor and is freely available for non-commercial users.

## 1 Introduction

Unlike genomes, changes in epigenomes are dynamic and reversible. This plasticity has been increasingly highlighted over a recent decade, since the spatiotemporal dynamics of epigenetic modification are shown to form the basis of diverse cellular processes such as cell differentiation and senescence. The variability of epigenomes is the main mechanism that cells acquire diverse and precise functions throughout the whole body of an individual to sustain the life of an organism. Furthermore, the dysregulation of the epigenetic variability has shown great biological implication for various cancer types, as it increases the adaptive potential of a cancer cell population against treatments (Mazor *et al*., 2016).

Among the diverse manifestations of epigenetic variabilities, cell-to-cell variability of DNA methylation states is one of the most actively investigated research topics. To measure the diversity of bulk cell population in terms of DNA methylation, it is necessary to identify the epigenetic configurations of individual cells. Single-cell bisulfite sequencing can directly resolve this challenge, but its cost makes it hardly applicable to a large-scale study. An effective alternative is to extract DNA methylation states co-occurring in a single sequencing read, and consider each pattern as a pseudo-barcode that identifies each cell. By measuring the diversity of DNA methylation patterns aligned at each genomic region, we can obtain a partial estimate of the true epigenetic diversity across cell population.

Despite many proof-of-concept experiments underscoring the utility of those DNA methylation heterogeneity measures in physiopathological conditions, a highly efficient toolkit for quantifying the extent of the heterogeneity is still lacking. To facilitate a large-scale functional study of DNA methylation heterogeneity based on bisulfite sequencing data, we developed a fast and lightweight software called *Metheor*. Here we present the functionality of Metheor and benchmark its performance against existing code implementations for DNA methylation heterogeneity measures. We believe this software will serve as a convenient toolkit helping researchers fully utilize the phasing information of DNA methylation states that have not been investigated actively so far.

## 2 Methods

### 2.1 Functionality and implementation

Given a bisulfite read alignment file, Metheor can compute all of the six DNA methylation heterogeneity measures proposed to date (Xie *et al*. (2011); Landau *et al*. (2014); Landan *et al*. (2012); Guo *et al*. (2017); Scherer *et al*. (2020); Figure 1A and B), and an additionally developed measure named local pairwise methylation discordance (LPMD). The detailed description for each of the measures are available in the Supplementary Information.

**Fig. 1.**
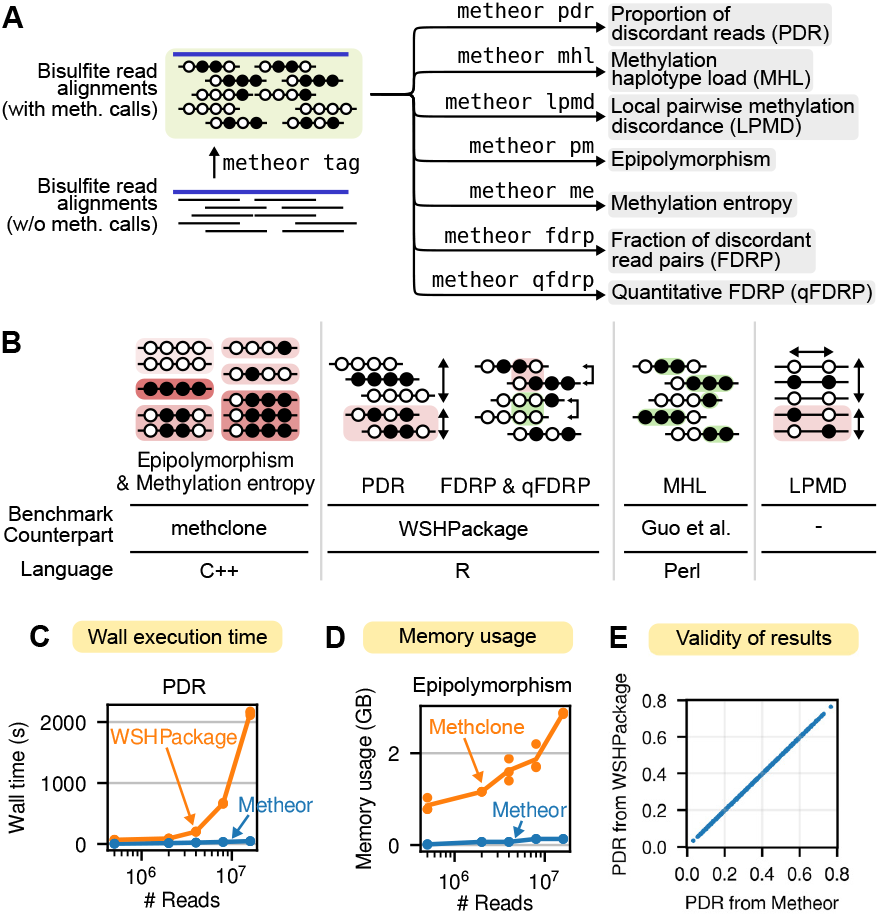
Overview of Metheor. (A) The input for Metheor is bisulfite read alignments tagged with Bismark methylation call strings. Using each of the seven subcommands shown, Metheor computes the corresponding DNA methylation heterogeneity measure. If reads were aligned with a tool other than Bismark, Metheor can still add tag for methylation call string with `metheor tag` subcommand. (B) Schematic diagram for DNA methylation heterogeneity measures and benchmark settings in this study. Guo et al. denote the Perl script provided by the authors along with the article proposing the utility of MHL. (C) Comparison of wall execution time required to compute PDR using WSHPackage and Metheor. (D) Comparison of memory usage during the computation of epipolymorphism using Methclone and Metheor. (E) Comparison of the PDR values computed by Metheor and WSHPackage.

The main algorithmic advantage of Metheor is that it is a “one-sweep” algorithm that iterates through the entire sequencing reads only once. On the other hand, a benchmark counterpart R package, WSHPackage, iterates through a set of CpGs specified by a user and fetches the reads covering each CpG using indexed alignment file. As the most time-consuming operation in this setting is fetching sequencing reads, we can estimate the time complexity of the methods using the total number of sequencing read accesses. For Metheor, the estimation is trivial; it requires *n* sequencing read accesses in total, where *n* is the number of aligned reads. For WSHPackage, the number of sequencing read accesses is *λ*×*n*, where *λ* is the average number of CpGs within a single sequencing read. Therefore, the performance advantage of Metheor compared to WSH will be determined by the coefficient *λ*. Notably, the empirical distribution of CpGs throughout the genome is uneven and known to form CpG-dense regions including CpG-islands. Reduced representation bisulfite sequencing (RRBS) predominantly targets those CpG-dense regions, so the coefficient *λ* for RRBS experiment is generally greater than 1 (Supplementary Figure 1). This is why the read-centric algorithm used in Metheor empirically runs faster than CpG-centric ones. Metheor also supports alignments generated not only by Bismark, but also other widely used methylation-aware aligners, by providing a subcommand to attach a tag denoting the methylation states of cytosines to each aligned read. This warrants the wide applicability of Metheor at the downstream of various bisulfite read processing pipelines. Metheor is implemented in Rust language and distributed via the conda package manager.

### 2.2 Performance benchmark

We compared the performance of our method with existing methods (methclone (Li *et al*., 2014), WSHPackage (Scherer *et al*., 2020) and perl scripts from Guo *et al*. (2017) for the six existing measures. Three different types of simulated sequencing datasets with varying numbers of reads were used (Supplementary Information). In brief, we generated two RRBS datasets (by simulation and real-world data subsampling) and pseudo-whole genome bisulfite sequencing experiments reproducing the methylation erosion scenario as proposed in Scherer *et al*. (2020). All the benchmark experiments were done on a server with Intel(R) Xeon(R) E7-4850 2.10 GHz CPU and 512GB of RAM.

## 3 Results

### 3.1 Execution time and memory usage benchmarks

We observed significant speedups (up to 300-fold depending on measure and data size) for the computation of all DNA methylation heterogeneity measures (Figure 1C, Supplementary Figure 2-4A). It is worth noting that the total running time of Metheor and WSHPackage were both linearly proportional to the number of sequencing reads in the input data. Notably, we could achieve extreme speedup for FDRP and qFDRP calculation using reservoir sampling-based approach (Supplementary Information). Since the utility of FDRP and qFDRP has been limited by the slow computation, we expect that Metheor will facilitate the use of those measures in further experimental setups. We also observed significant reduction in memory footprints (Figure 1D, Supplementary Figure 2-4B).

WSHPackage allows PDR, FDRP and qFDRP to be calculated using multiple threads. To examine whether it can achieve performance comparable to Metheor when multiple threads are used, we measured the total running time for PDR, FDRP and qFDRP using up to 32 threads. Surprisingly, we found that Metheor, with only a single thread, showed remarkable performance improvement (13.7∼175.8-fold faster) even compared to WSHPackage using 32 threads.

### 3.2 Validity of the result

We verified that the methylation heterogeneity levels computed by Metheor are identical to the results from the reference implementations at both individual CpG or epiallele level (Figure 1E, Supplementary Figure 5). For sampling-based measures (FDRP and qFDRP), we could instead show that the results were highly correlated. Altogether, these results ensure the reliability of the results from Metheor throughout all the measures it supports.

## Supplementary Material for

### 1 Supplementary information

In this supplementary information, we describe both the details of the algorithms used in Metheor implementation and simulated data preparation used for benchmark.

#### 1.1 Data structure and notations

Metheor parses bisulfite alignment files using `rust-htslib` library (Köster, 2016). While iterating through the aligned reads, we utilize a compact representation of aligned bisulfite reads to keep the memory requirement as minimum as possible. Throughout the discussion, we denote an aligned bisulfite read as *r*, which is internally represented as a composite data structure consisting of the following member variables:

- `start_pos`: The (0-based) leftmost position with regard to the reference genome where the read was aligned. Its value is accessed by a method getStartPos.
- `end_pos`: The (0-based) rightmost position with regard to the reference genome where the read was aligned. Its value is accessed by a method getEndPos.
- `cpgs`: A vector of CpGs covered by the read. The number of CpGs is accessed by a method getNumCpgs, and the position of the first (leftmost) CpG is accessed by a method getFirstCpgPosition.

Individual CpG on the sequencing read is also represented by a dedicated data structure with the following member variables:

- `abspos`: The (0-based) absolute position of the CpG with regard to the reference genome (i.e., corresponding chromosome). Its value is accessed by a method getPosition.
- `relpos`: The (0-based) relative position of CpG within the sequencing read. Maintaining this variable facilitates simple operations for CpGs within a read.
- `is methylated`: A boolean value denoting whether the CpG is determined to be methylated in the read. Its value is accessed by a method IsMethylated.

The methylation state of each CpG can be easily determined by directly parsing the Bismark methylation string (BAM tag specified as ‘XM’) (Krueger and Andrews, 2011). Therefore, it is required for an input BAM file to have Bismark methylation strings in order to run Metheor. For those who prefer methylation-aware aligners other than Bismark, we provide an auxiliary command named `tag` to add XM-tag automatically to any BAM file.

In the following sections, we discuss the biological motivations of each methylation heterogeneity measure and algorithmic details for the computation of them in Metheor.

#### 1.2 Computation of proportion of discordant reads (PDR)

Due to the processive nature of the catalysis driven by DNA methyltransferases and demethylases, local DNA methylation states are correlated with each other. In other words, it is highly likely to observe a methylated CpG when the nearby CpGs are methylated, and vice versa. In cancers, this local homogeneity of DNA methylation states is known to be eroded and is also known to be associated with the clinical outcome of a patient. In this regard, PDR measures how much the local homogeneity of DNA methylation states is eroded in a given genomic region. In the definition of PDR (Landau *et al*., 2014), a read is defined as *discordant* if CpGs covered by the read has both methylated and unmethylated states (representing *locally disordered* methylation states), otherwise a read is defined as *concordant*. PDR is a CpG-wise measure, that is, a single PDR value is assigned for a CpG. For a set of reads covering the CpG, PDR is defined as a proportion of discordant reads. For example, a CpG covered by reads harboring consistent methylation states (either fully-methylated or fully-unmethylated) has PDR of 0, and otherwise has PDR greater than 0.

In this section, we describe how the computation of PDR is implemented in Metheor using a **single sweep** of the given alignment file. As a result, a hashmap associating each CpG to a corresponding PDR value is obtained by Algorithm 1. While iterating bisulfite reads in the given alignment file, we maintain a temporary hashmap *T* linking each CpG to a number of associated concordant/discordant reads or *n*_*c*_ and *n*_*d*_. For each read *r* and for each CpG within the read, *n*_*c*_ and *n*_*d*_ in *T* is continuously updated based on the concordance state of read *r*, which is determined by IsDiscordant method (Algorithm 2). Note that by assuming that the alignment file is sorted by genomic coordinates in an increasing order, it can be ensured that a CpG will not be processed any more when we encounter a read whose leftmost CpG position is greater than that CpG. By finalizing those CpGs to the hashmap *H*, we can reduce the amount of working set in *T*.

##### Algorithm 1

Computation of proportion of discordant reads (PDR)

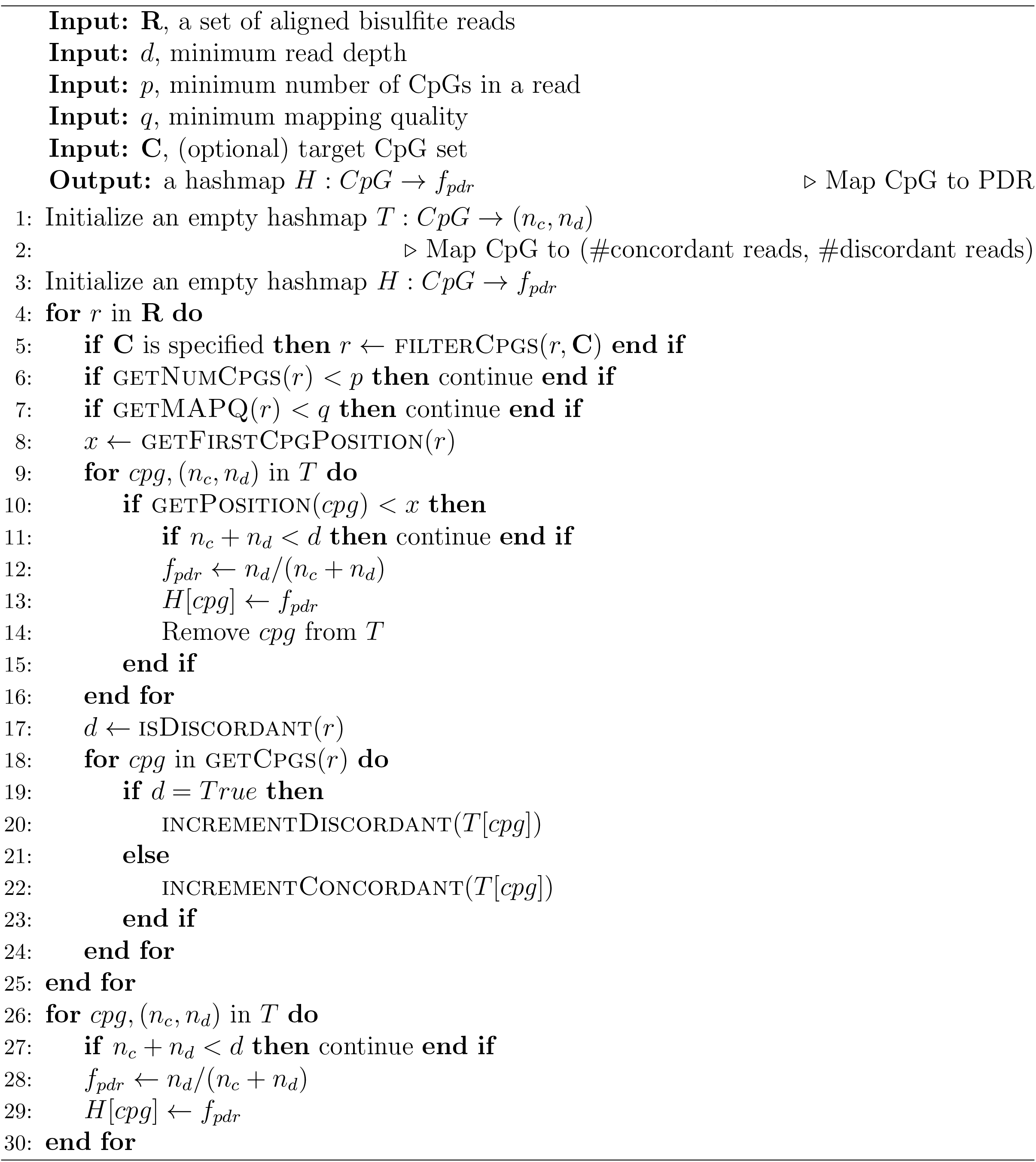

##### Algorithm 2

Determining whether a bisulfite read is discordant

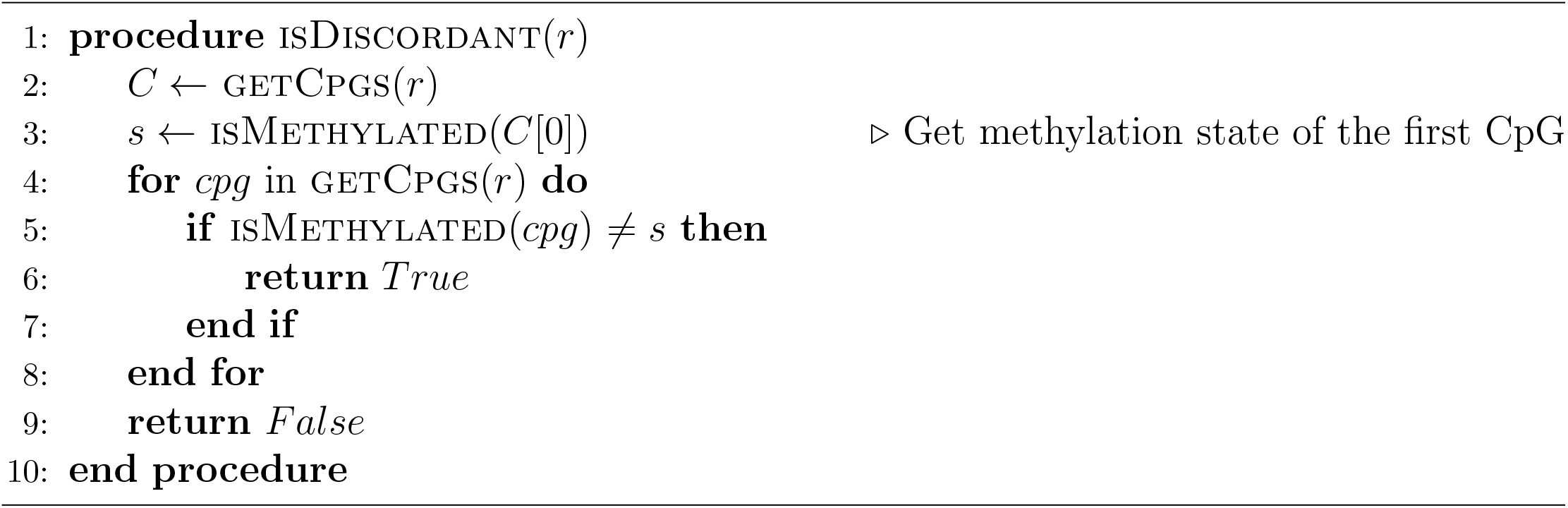

#### 1.3 Computation of local pairwise methylation discordance (LPMD)

In this study, we introduce LPMD as a new measure to quantify the local concordance of DNA methylation states. The conceptual basis of LPMD is similar to PDR as they both are aware of local homogeneity of DNA methylation states, but they are different in that LPMD explicitly takes the distance between CpGs into consideration. In detail, LPMD is defined as a fraction of CpG pairs within a given range of genomic distance (i.e., CpG pairs more distant than *m* bp and closer than *M* bp) and therefore, LPMD is defined for a *pair* of CpGs, but not for a single CpG. Importantly, we note that LPMD does not depend on length of sequencing read. Considering that there is an increased tendency of observing discordant reads solely by chance based on definition in PDR, LPMD proves to be a novel method for measuring DNA methylation heterogeneity.

The goal of Algorithm 3 is to obtain a hashmap associating a pair of CpG to a corresponding LPMD value. This algorithm also computes LPMD value for each pair of CpGs after a **single sweep** of the given alignment file. Extracting all pairwise concordance/discordance statistics is performed using ComputePairwiseStats method (Algorithm 4), which is a variation of two-pointer algorithm that keeps the minimal set of CpGs needed for further pairwise examinations.

##### Algorithm 3

Computation of local pairwise methylation discordance (LPMD)

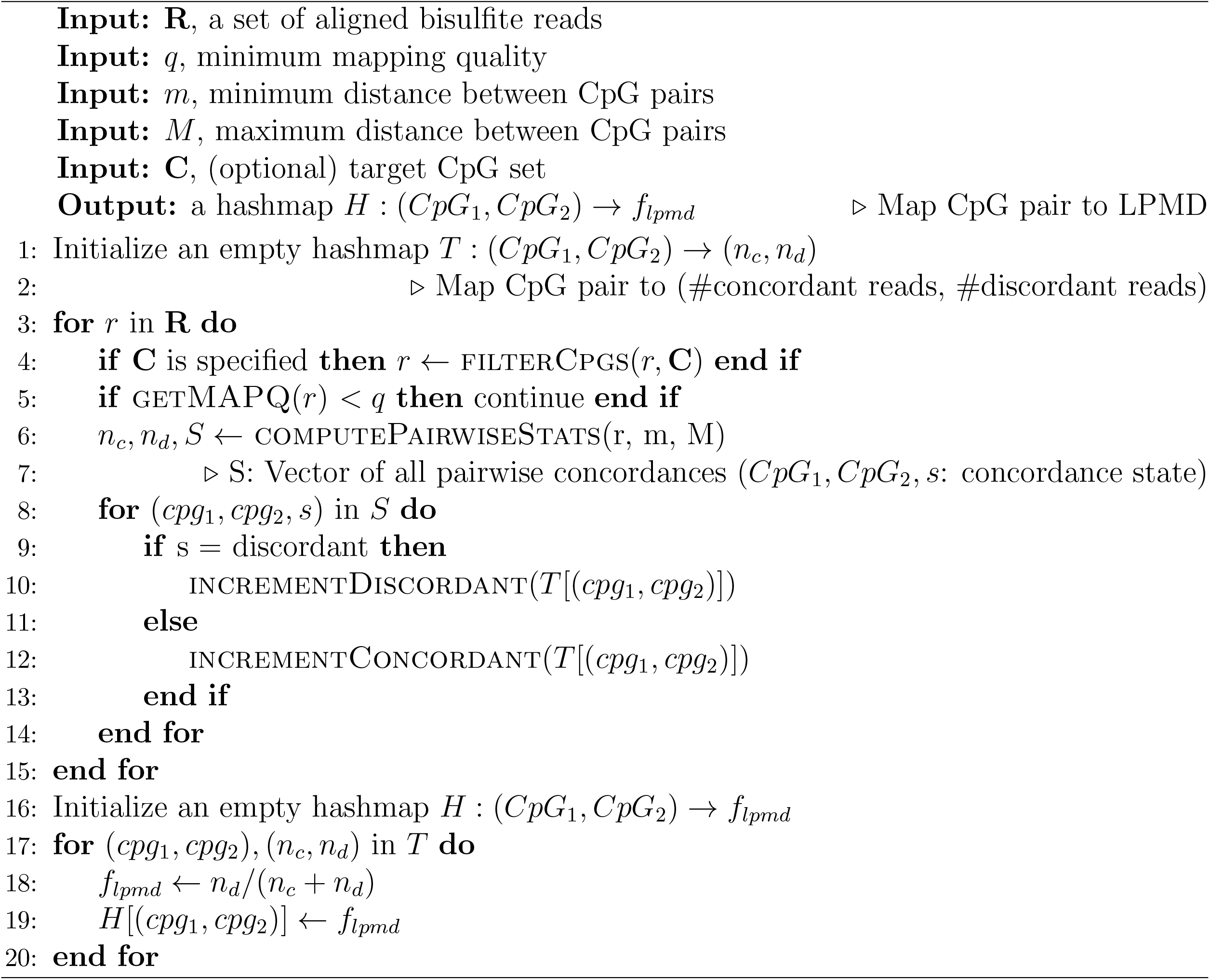

##### Algorithm 4

Extracting all pairwise concordance/discordance statistics from a read

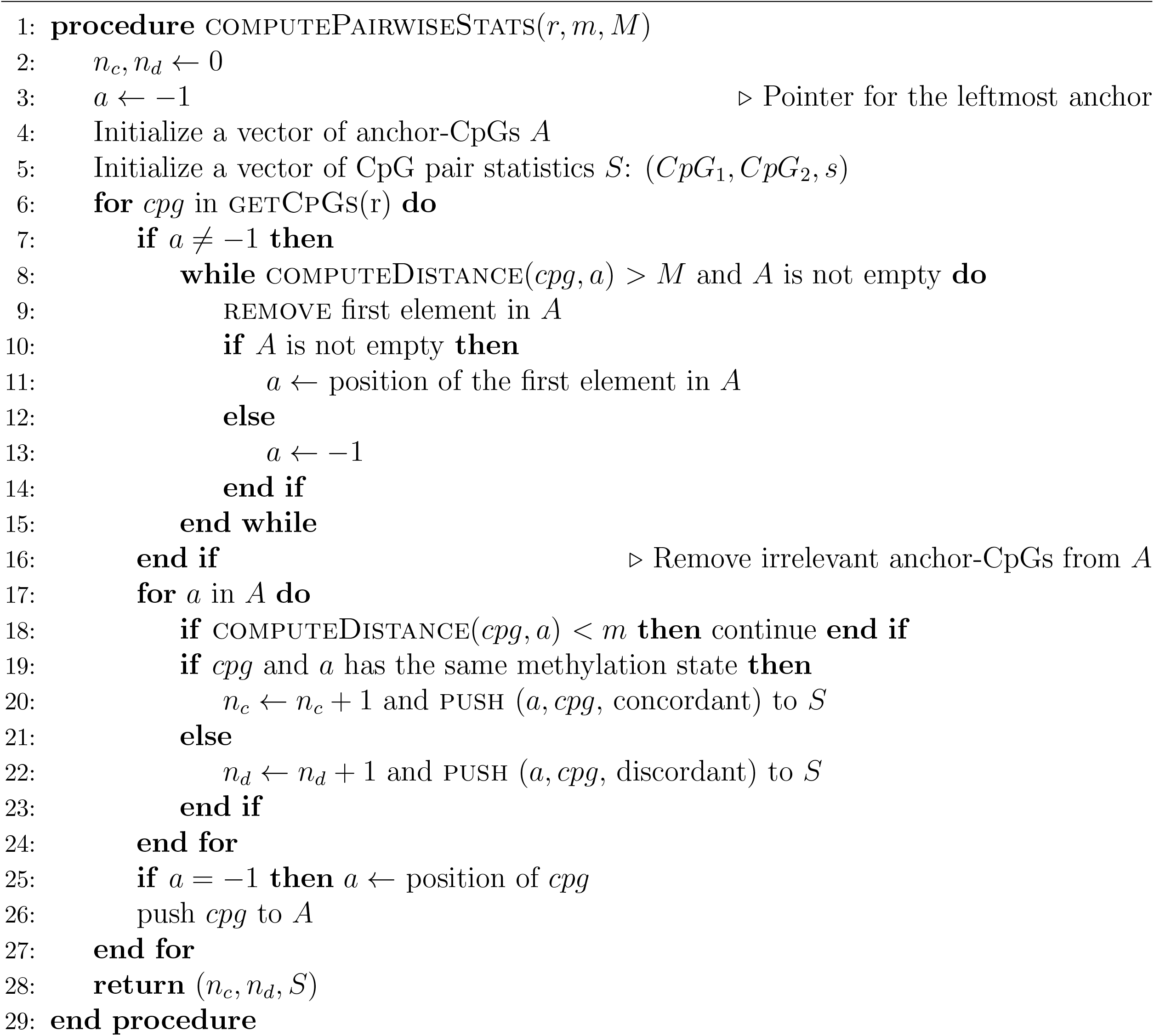

#### 1.4 Computation of methylation haplotype load (MHL)

The concept of MHL is also based on the local homogeneity of DNA methylation states, or co-methylation, due to the processivity of enzymes responsible for methylation and demethylation of cytosines (Guo *et al*., 2017). While PDR and LPMD focus on how much the tendency of co-methylation is perturbed in the given population of cells, MHL focuses on how well the methylation haplotypes (i.e., *stretch* of consecutive methylated CpGs) are conserved throughout the cell population for a given genomic region. The basic concept of MHL is to systematically identify the genomic blocks harboring CpGs with tightly coupled methylation states. In detail, MHL is computed as a fraction of observed *fully methylated stretches* out of the all stretches of every possible lengths. Notably, the authors proposed that giving more weights to longer methylated stretches heuristically worked well. Altogether, an intuitive definition of MHL can be formulated as below.

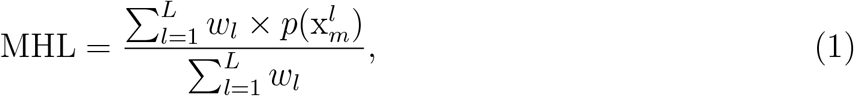

where 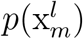 denotes the fraction of fully methylated stretches of length *l* out of all possible stretches, and *w*_*l*_ represents the weight given to a fraction of methylated stretch of length *l*. In the implementation of Metheor, we sticked to *w*_*l*_ = *l* according to the suggestion of the original authors (Guo *et al*., 2017).

In the current version of Metheor implementation, a hashmap associating each CpG to a corresponding MHL value is obtained by Algorithm 5 with a **single sweep** of read alignments. For an efficient computation of MHL, we maintain a composite data structure named `AssociatedReads` that contains the summary statistics of methylated stretches associated with each CpG. `AssociatedReads` consists of the following member variables:

- `pos`: Position of CpG of interest for which this object summarizes the information of associated reads.
- `stretch_info`: Hashmap *S* : *l* → *n*_*l*_, which represents the count *n*_*l*_ of methylated stretch of length *l*.
- `num_cpgs`: Vector of CpG counts for all the associated reads. Accessed by a method getNumCpgs.
- `max_num_cpgs`: max(`num cpgs`). Accessed by a method getMaxNumCpgs.

In particular, introducing a hashmap `stretch_info` is the key idea for the efficient computation of MHL. Using methylation states of CpGs covered by a read, we can compute a read-wise statistics of methylated stretch as Algorithm 6, and using `stretch_info`, we can easily compute MHL values using Algorithm 7.

##### Algorithm 5

Computation of methylation haplotype load (MHL)

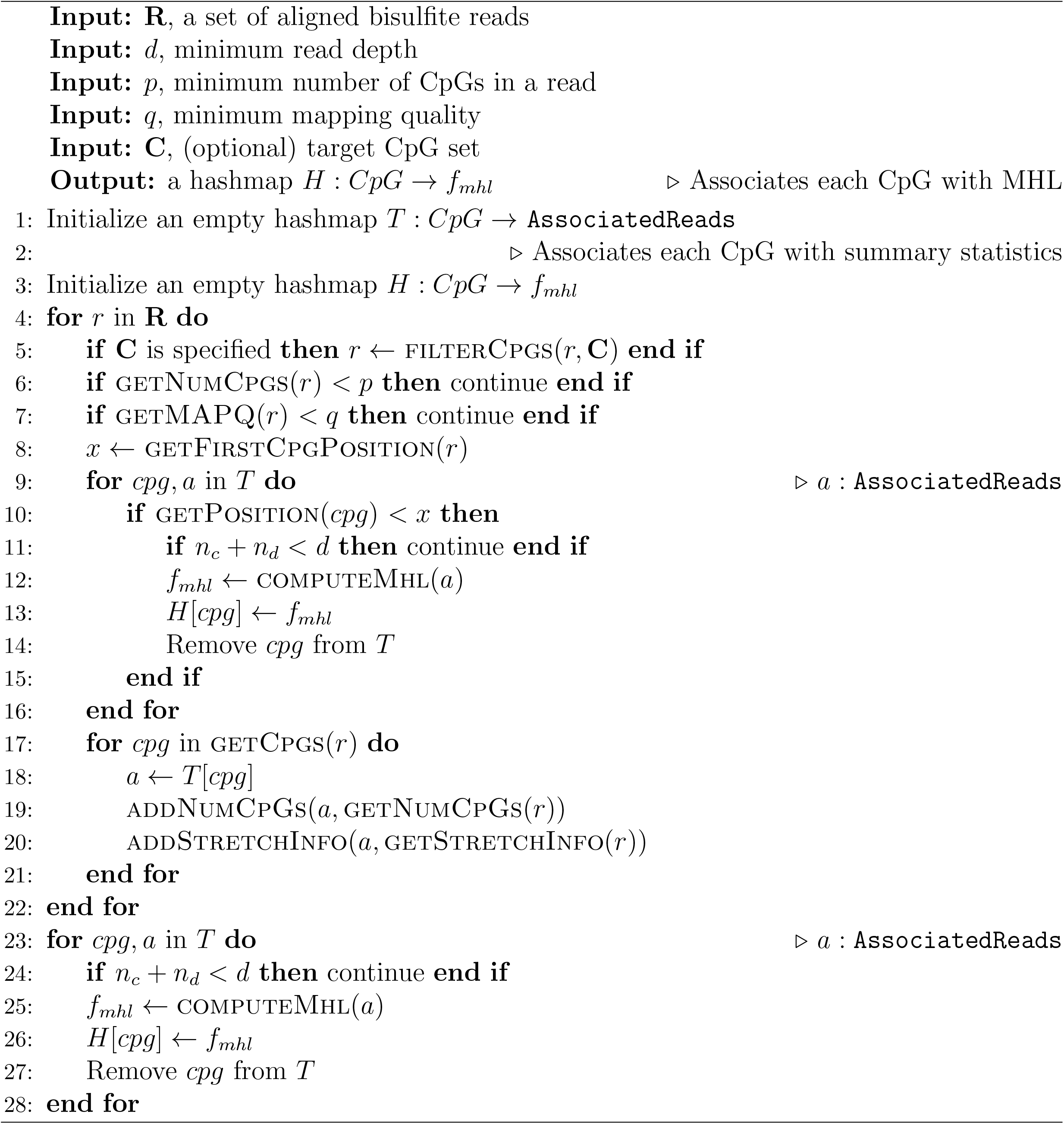

##### Algorithm 6

Extracting metylation stretch_information from a read

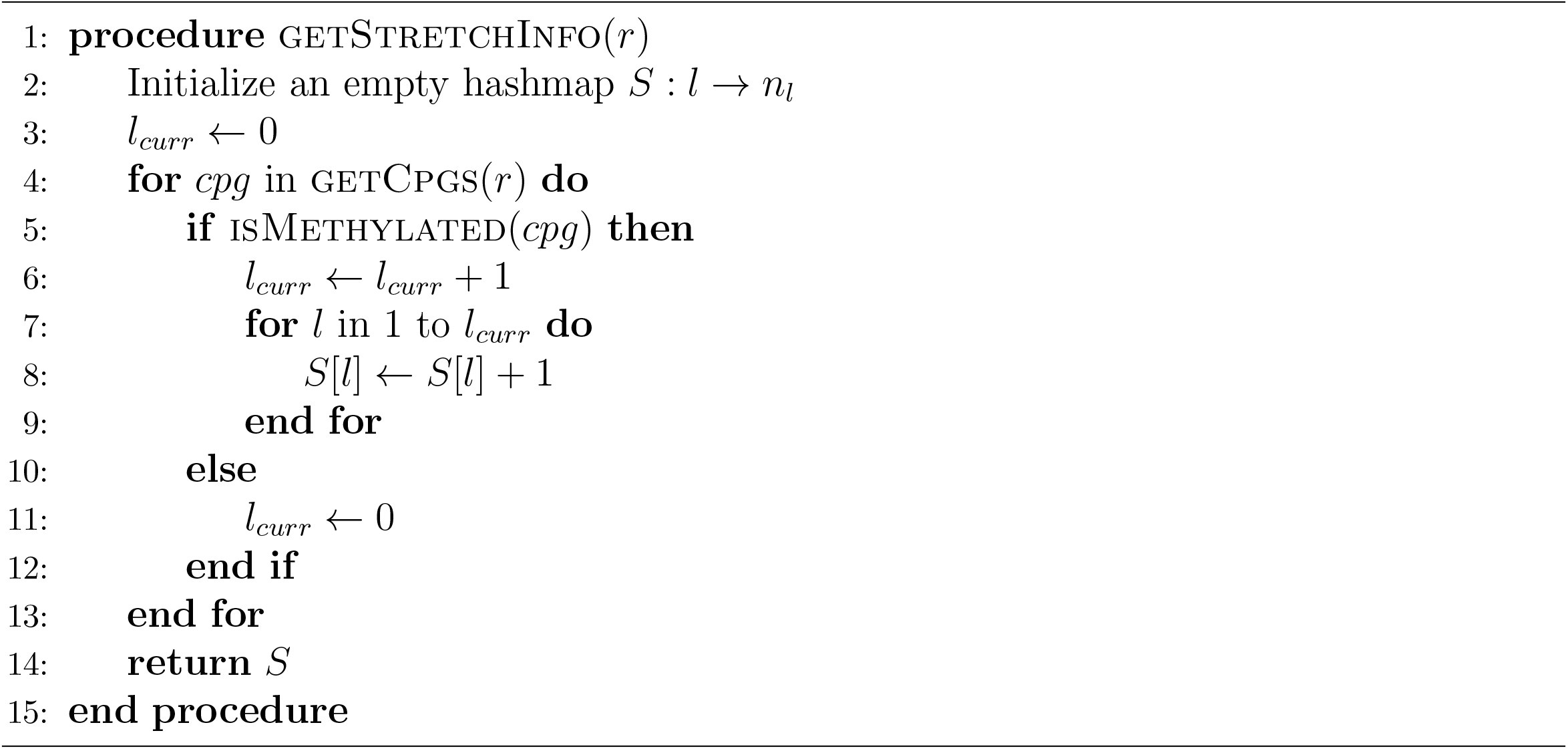

##### Algorithm 7

Computing MHL from summary statistics of associated reads

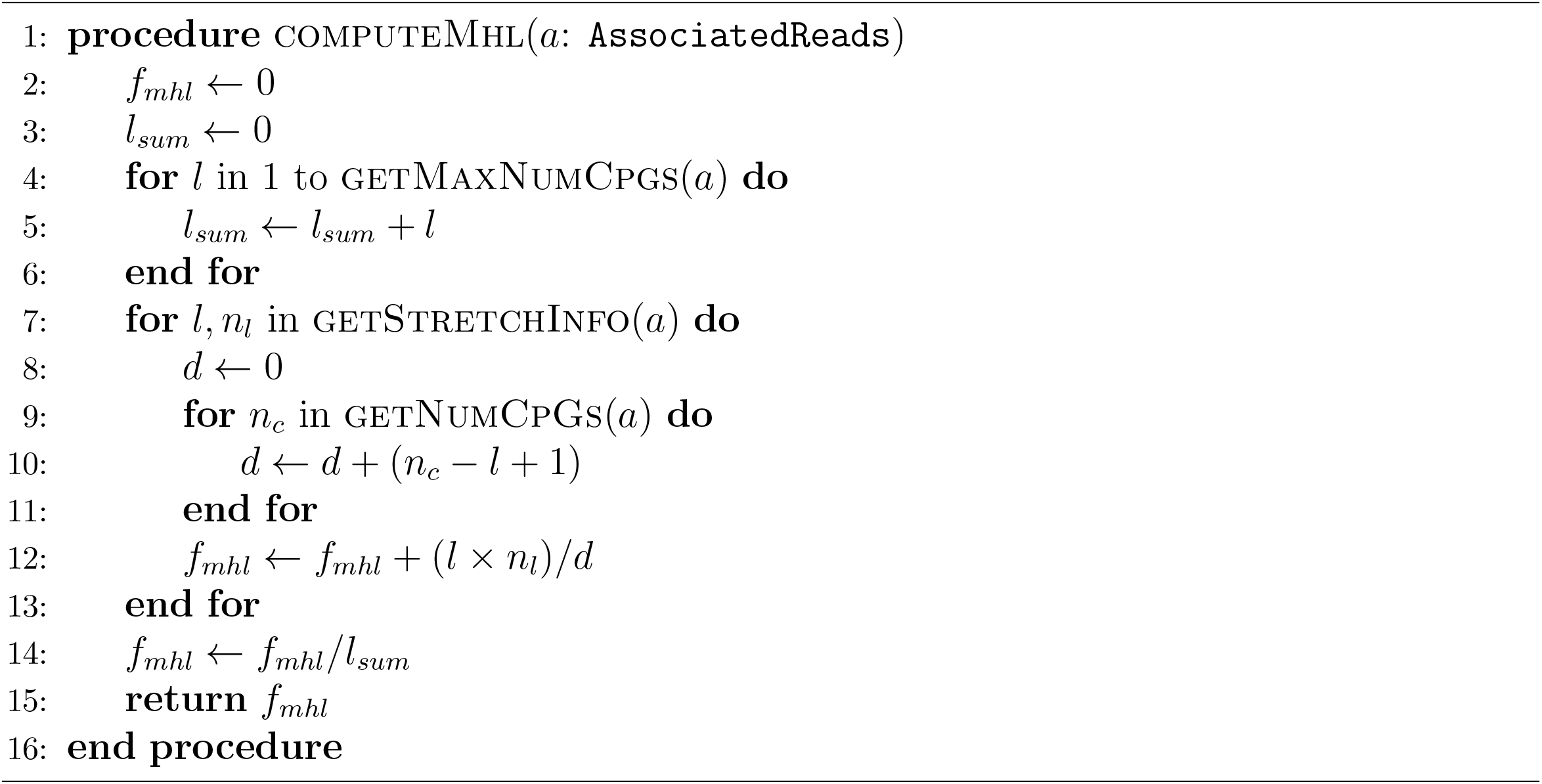

#### 1.5 Computation of epipolymorphism (PM) and methylation entropy (ME)

PM (Landan *et al*., 2012) and ME (Xie *et al*., 2011) are closely related measures that are defined for phased methylation states, or epialleles, so we discuss these two measures together in this section. The goal of these measures is to quantify the diversity of DNA methylation states of a given cell population. However, identifying the whole phased DNA methylation states of each cell to compute and calculating their diversity is not feasible. To circumvent this problem practically, PM and ME measure the diversity of the *methylation patterns*, or epialleles, formed by four consecutive CpGs covered by a single. bisulfite read. There are 2^4^ = 16 possible methylation patterns in total. Note that the concordance/discordance of the methylation states of nearby CpGs are not of interest to PM and ME, but they are only interested in how diverse those 16 methylation patterns are. To distinguish the four consecutive CpGs themselves from the methylation states of them, we term the former as *CpG quartet* and the latter as *methylation patterns* or *epialleles*.

The following algorithms (Algorithm 8 and 9) show how we obtain a hashmap *H* associating each CpG quartet with corresponding PM and ME values through a **single sweep** of the alignment file.

##### Algorithm 8

Computation of epipolymorphism (PM)

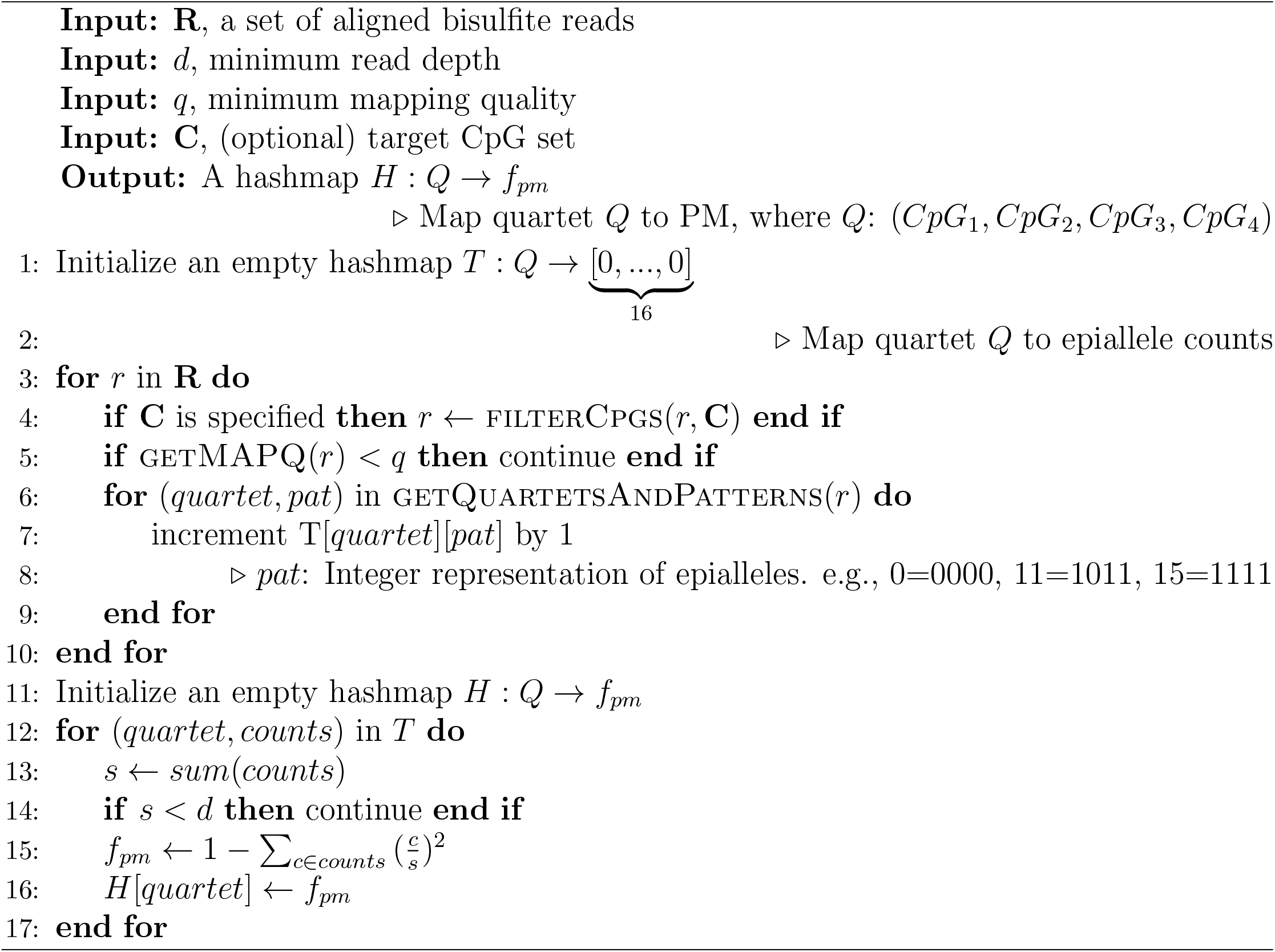

##### Algorithm 9

Computation of methylation entropy (ME)

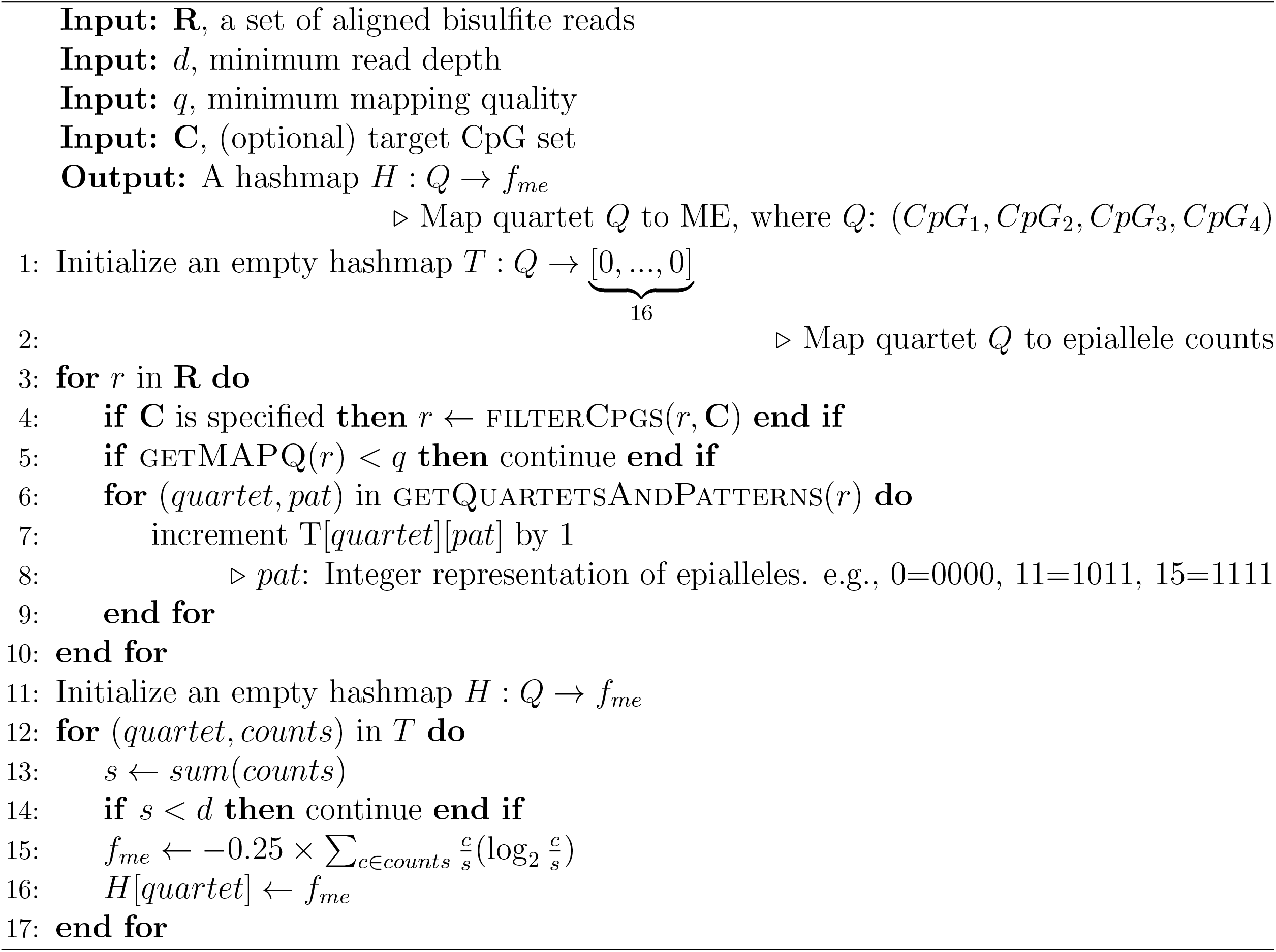

#### 1.6 Computation of fraction of discordant read pairs (FDRP) and quantitative FDRP (qFDRP)

FDRP and qFDRP are measures of epigenetic diversity within a cell population that were first proposed in Scherer *et al*. (2020). While PM and ME quantify the epiallelic diversity in terms of CpG quartets, FDRP and qFDRP allow the computation of epiallelic diversity in a single CpG resolution. The key principle underlying the FDRP and qFDRP is as follows. When epialleles are perfectly homogeneous for a short genomic region, any two sequencing reads aligned to that region will have identical methylation states for CpGs that are common to the two reads. On the other hand, as epialleles become more diverse, it is more likely to observe a read pair that have different methylation states for common CpGs. Based on this notion, FDRP and qFDRP compute a CpG-wise epigenetic diversity by examining pairs of sequencing reads covering the CpG. Since the time required for all pairwise examination of sequencing reads increases exponentially along with the sequencing depth, the authors adopt a read sampling strategy to make those measures be computed in a feasible time. Therefore, the maximum number of sampled reads, or *M*, is a crucial parameter modulating the balance between the precision of the measures and the computing time.

Algorithm 10 and 11 show how Metheor obtains a hashmap associating each CpG with FDRP and qFDRP values. Like all the other measures, Metheor computes FDRP and qFDRP using a **single sweep** of sequencing reads. Similarly to MHL, FDRP and qFDRP require their own `AssociatedReads` data structure tailored for the efficient calculation of the measures.

- `pos`: Position of CpG of interest for which this object summarizes the information of associated reads.
- `reads`: Vector of sequencing reads. Maximum length of this vector is kept to `max depth`
- by reservoir sampling.
- `num_total_read`: Number of the reads actually covering the CpG of interest.
- `num_sampled_read`: Number of reads in the current set of sampled reads.
- `max_depth`: Maximum number of reads allowed for sampled reads.

To reduce the memory usage during iteration and perform read sampling efficiently, Metheor utilizes a reservoir sampling when the number of reads covering the CpG exceeds `max_depth` or parameter *D*. For convenience, we will denote `num_sampled read` as *n* in this section. When the CpG is ready to be finalized (i.e., it is guaranteed that no more reads cover the CpG), we examine all the *n*(*n* − 1)*/*2 read pairs and determine whether the pair is discordant (i.e., at least one CpG common to the two reads have different methylation state) or concordant (i.e., all CpGs common to the two reads have identical methylation states) to compute FDRP. In the case of qFDRP, the normalized hamming distance (i.e., the number of CpGs with different methylation states divided by the number of common CpGs) is used instead. Finally, we obtain FDRP by dividing the number of discordant read pairs by *n*(*n* − 1)*/*2, and we obtain qFDRP by dividing the sum of normalized hamming distance also by *n*(*n* − 1)*/*2.

##### Algorithm 10

Computation of fraction of discordant read pairs (FDRP)

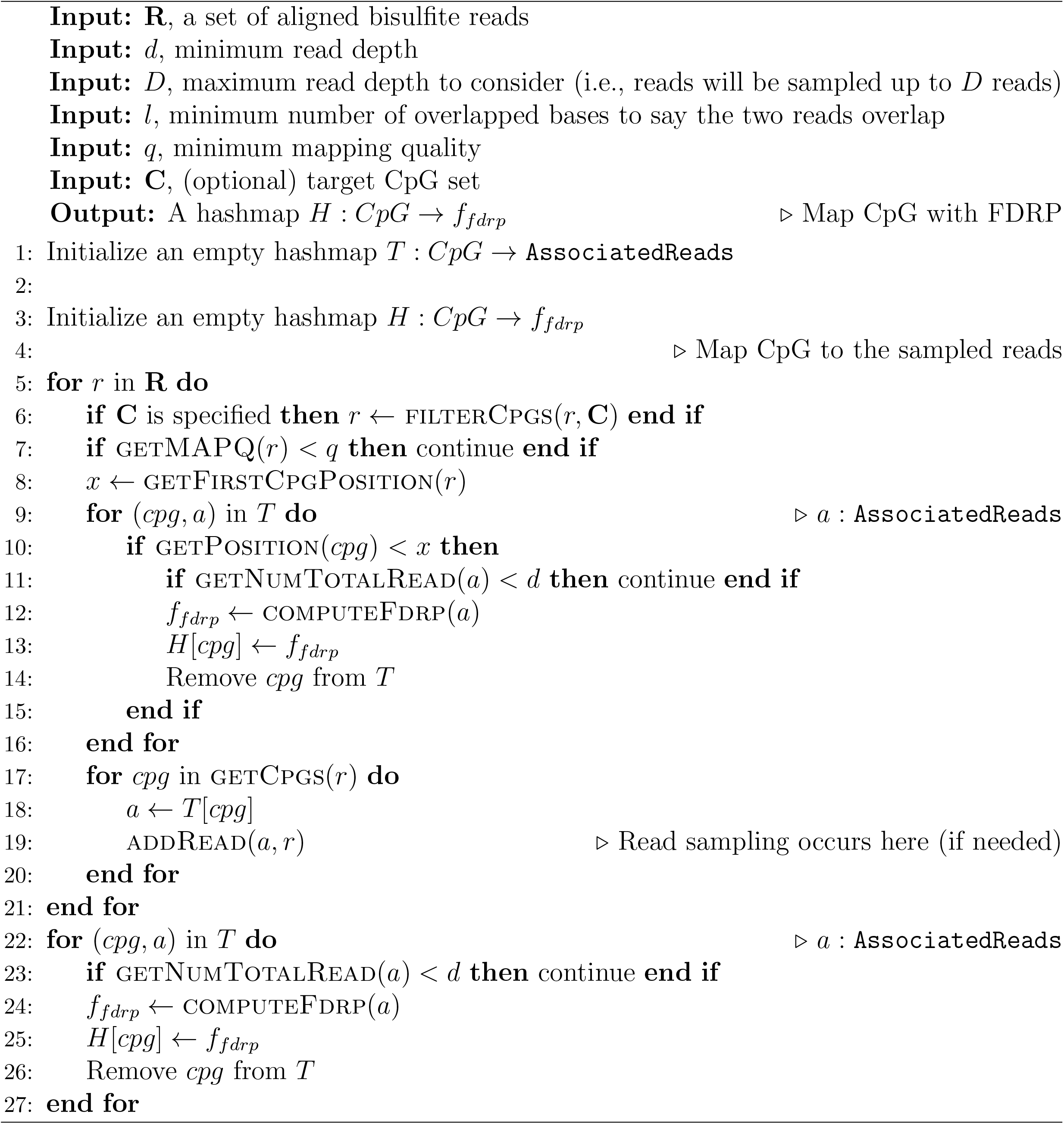

##### Algorithm 11

Computation of quantitative FDRP (qFDRP)

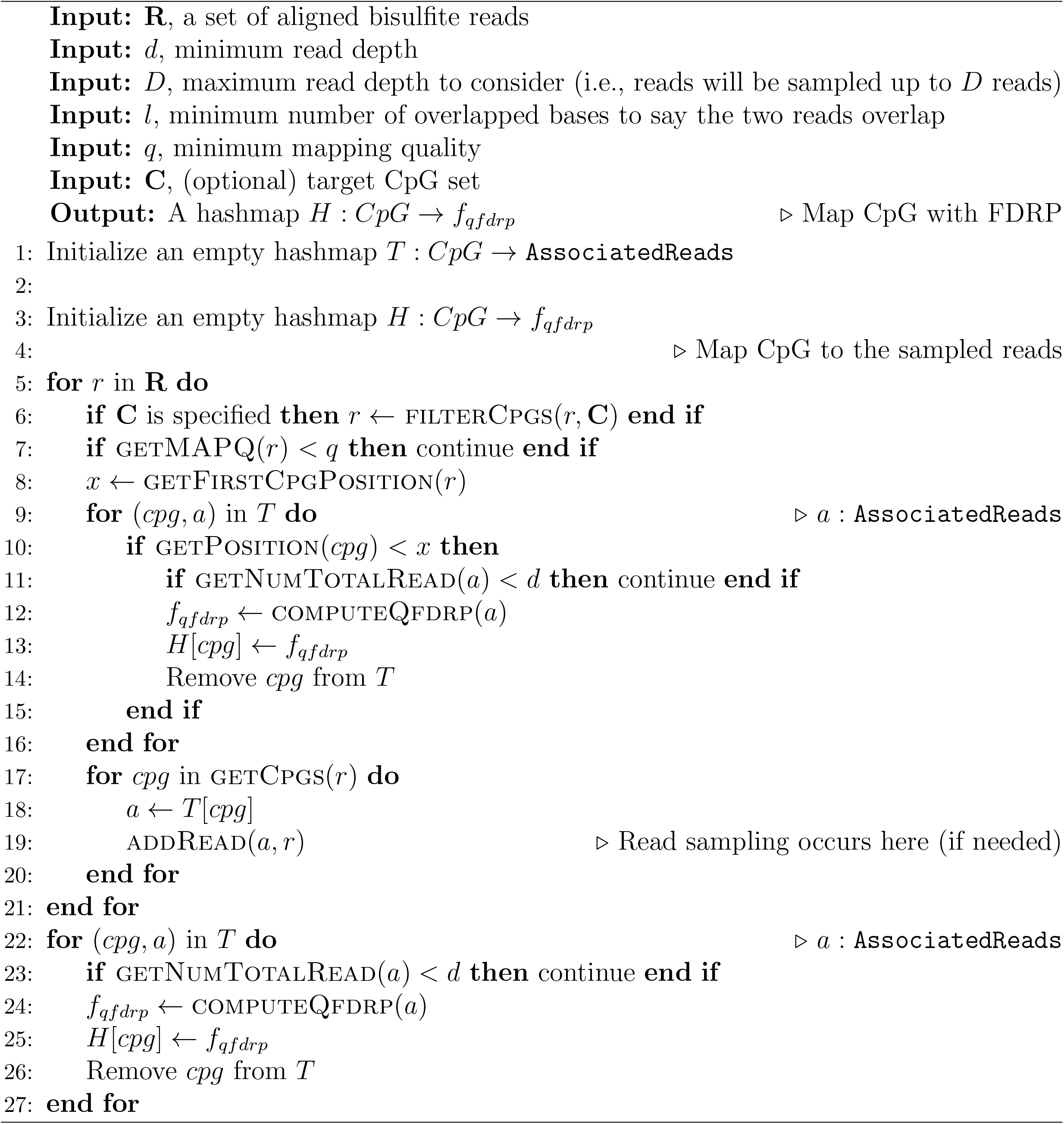

**Supplementary Fig. 1.**
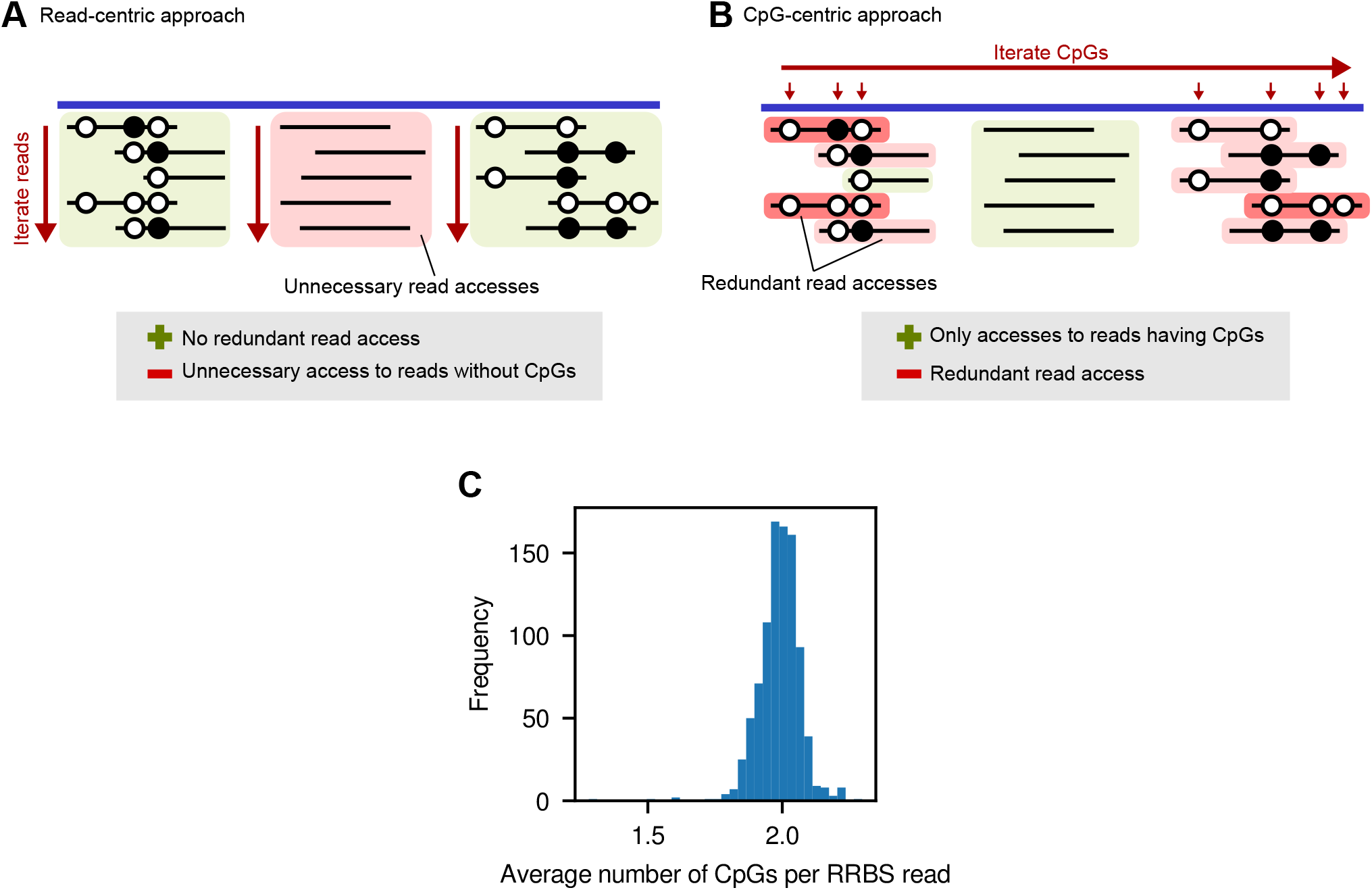
Algorithmic advantages of Metheor. (A) Schematic illustration of read-centric approach. This approach iterates through aligned reads, as illustrated by red arrows. Therefore, it is guaranteed that only one read access occurs for each read. However, since we cannot determine whether the read covers any CpG or not before accessing the read, there inevitably are unnecessary accesses to reads that do not cover CpG. (B) Schematic illustration of CpG-centric approach. This approach iterates through an externally specified set of CpGs. Therefore, reads that do not cover CpGs can be excluded from the computation, but redundant read accesses occur because there are reads covering more than one CpGs. (C) Average numbers of CpGs per single RRBS read (*λ*) computed for each of 928 RRBS experiments on CCLE cell lines.

#### 1.7 Insights into the algorithmic advantages of Metheor

The main algorithmic advantage of Metheor comes from the fact that it only reads through the entire BAM file only once (Supplementary Figure 1A). In this section, we call it as *read-centric* approach, as each sequencing read is processed only once. On the other hand, existing methods for the computation of PDR, FDRP, qFDRP (by WSHPackage) and MHL (by the Perl script provided by the authors) takes *CpG-centric* approach that iterates through each CpG specified by a user, and fetches the reads covering that CpG using BAM index (Supplementary Figure 1B). Even though the alignment index allows the access of reads quickly, such read access consumes the largest portion of the running time in the execution of the program. In this context, we here provide a brief insight into the algorithmic advantages in Metheor along with our empirical observations supporting the discussion.

The overhead of read-centric approach is that reads with no CpGs are also accessed. On the other hand, the overhead of CpG-centric approach is that redundant read accesses are needed for reads with more than one CpGs. To numerically compare the effect of the two potential drawbacks, we first denote the total number of aligned reads as *n*. Then, the number of read accesses in read-centric approach is trivially guaranteed to be *n*. For CpG-centric approach, we can notice that the number of access for a read is exactly same to the number of CpGs covered by the read. Therefore, the total number of read access in CpG-centric approach is the total number of individual cytosines covered by sequencing reads. When we denote the average number of covered CpGs per read as *λ*, we can conclude that CpG-centric approach requires *λ × n* read accesses. Thus, the value of *λ* for a dataset decides which approach is favored over the other.

To assess the empirical value of *λ* in general RRBS experiments, we collected the statistics from a large number of 928 public RRBS experiments on CCLE cell lines (Ghandi *et al*., 2019). Raw RRBS sequencing reads were downloaded from SRA under study accession SRP186687, adapter-trimmed using trim-galore, and aligned to the reference genome with Bismark. As shown in Supplementary Figure 1C, we discovered that the values of *λ* is greater than 1 in every case, supporting that read-centric approach in realistic samples.

#### 1.8 RRBS read simulation

To benchmark the performance of Metheor, we simulated a realistic RRBS data with various numbers of reads according the the procedure described in this section. First, the whole hg38 reference genome was digested *in silico* by a virtual restriction enzyme MspI having restriction site 5’-C|CGG-3’. It produced 2,317,722 genomic fragments in total. To imitate the experimental size-selection procedure, we only kept the fragments with size range between 50-200bp, and obtained 531,112 fragments. Then, we generated a simulated RRBS dataset by sampling reads from those fragments. Nine different sequencing data were generated with the number of sampled reads configured as follows: (1) 500K, (2) 2M, (3) 4M, (4) 8M, (5) 16M, (6) 20M, (7) 50M, (8) 100M and (9) 200M. To simulate a sequencing read, we randomly sampled a restriction fragment with replacement and determined the methylation level *β* using distribution *Beta*(0.25, 0.25). For each CpG covered by the read, it was methylated with probability *β*, i.e., the cytosine is converted to thymine with probability 1 − *β*. At the same time, a Bismark XM-tag was also simulated for each read accordingly. The orientation of the read (forward/reverse) was randomly determined with probability of 0.5. For convenience, read mapping quality was fixed to 40, base quality was fixed to 41 (‘J’ in ASCII representation) and no sequencing errors were introduced. Finally, all the simulated aligned reads were written to BAM files.

#### 1.9 Pseudo-WGBS read simulation

To simulate pseudo-WGBS reads, we followed a read simulation strategy used in Scherer *et al*. (2020). We especially adopted a scenario imitating DNA methylation erosion in the short genomic stretches throughout the whole genome. The whole simulation procedure is described as follows. First, we selected *N* random genomic regions across the genome (except chromosome 22, X and Y) in order to reduce the computational load while keeping the sequenced region unbiased for the whole genome. Note that this is why we call this simulated dataset pseudo-WGBS, not WGBS. All regions had fixed size of 50kbp, and no regions were allowed to have bases other than A, C, G and T. Then, for each region, we randomly divided the region into three segments. Each segment was forced to be at least 150bp long. We subsequently sampled sequencing reads from first and last segments (which we call first and second subregion, respectively), and kept all the CpGs on the sampled reads fully methylated. In this process, the number of sequencing reads to be sampled from those two subregions was determined proportionally to length of each corresponding region. Specifically, this number was calculated as 50kbp *×* (length fraction of each subregion). On the other hand, the segment in the middle (which we call erosion subregion) was processed in more sophisticated way to simulate the process of DNA methylation erosion. First of all, to represent stochastic methylation erosion occurring during cell proliferation, we sampled a random number (`replicate_num`) in the range of [2,10], following Scherer *et al*. (2020). Since we should consider both the `replicate_num` and length fraction of erosion region in deciding total number of sequencing reads to be sampled from this subregion, we initially sampled small number of samples from erosion subregion and concatenated resulting FASTQ files for `replicate_num` times. Also, we added two random factors in sampling sequencing reads from erosion subregion. The first was deciding whether to erode the methylation states of CpGs within erosion subregion, and the second was to determine the level of methylation erosion. For the first factor, we randomly sampled a floating-point number in the range of [0,1] and eroded the erosion region if this random number was above 0.5, which represents erosion probability of 0.5. For the second factor, we randomly sampled an integer in the range of [1,100] and set this as level of methylation erosion using Sherman (https://www.bioinformatics.babraham.ac.uk/projects/sherman). After independently processing *N* regions as explained above, we concatenated all the resulting FASTQ files into a single FASTQ file to obtain the final simulated sequencing reads.

**Supplementary Fig. 2.**
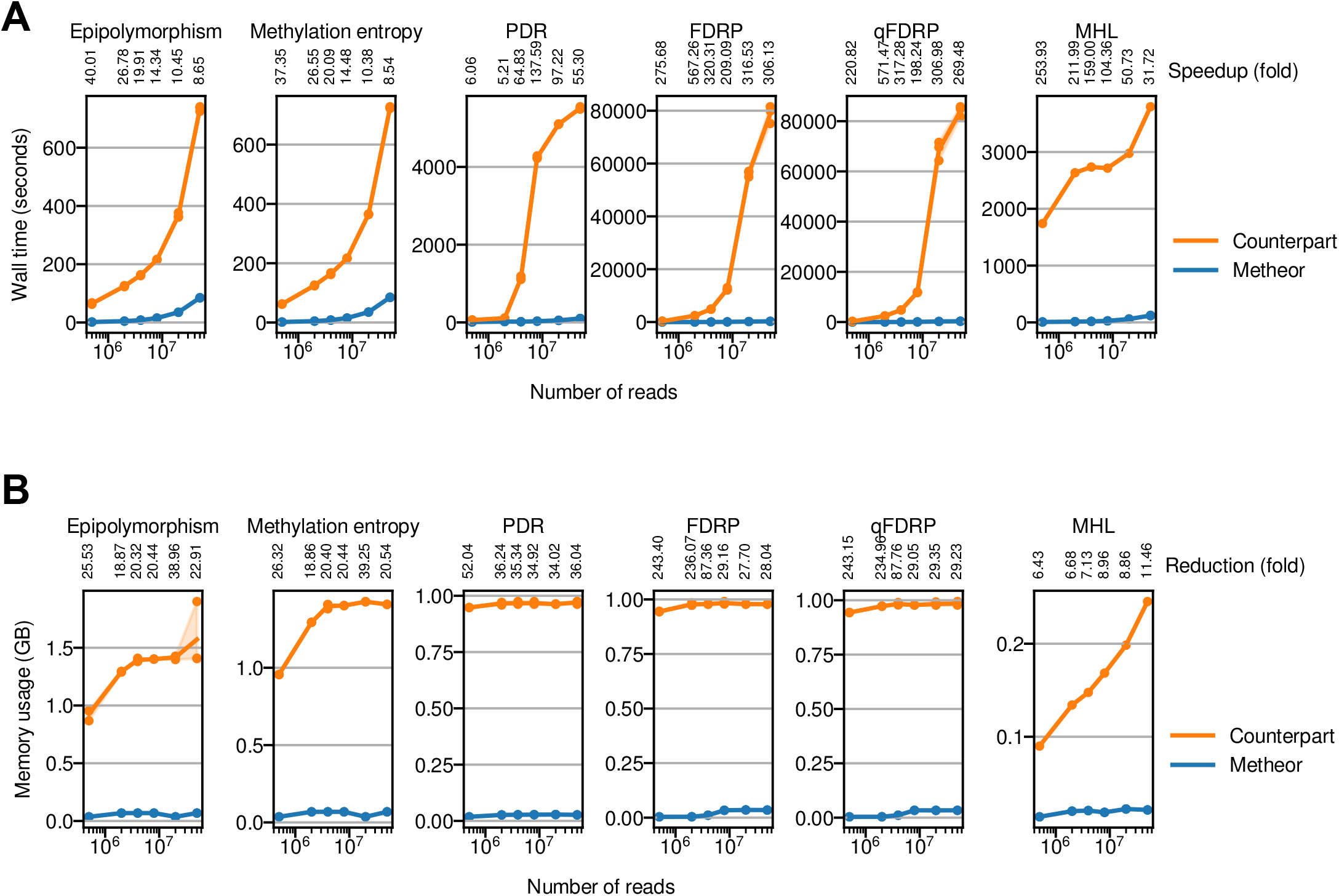
(A) Benchmarking the running time of Metheor using simulated RRBS dataset. Values below the name of each measure denote the amount of speedup (in fold) in Metheor compared to its benchmark counterpart. (B) Benchmarking the memory usage of Metheor using simulated RRBS dataset. Values below the name of each of the measures denote the amount of memory usage reduction (in fold) in Metheor compared to its benchmark counterpart. All the benchmarking experiments were repeated for three times, except for MHL. Lines denote the average wall time and shades represent the 95% confidence interval. Wall time and memory footprint for MHL computation was measured for only once.

**Supplementary Fig. 3.**
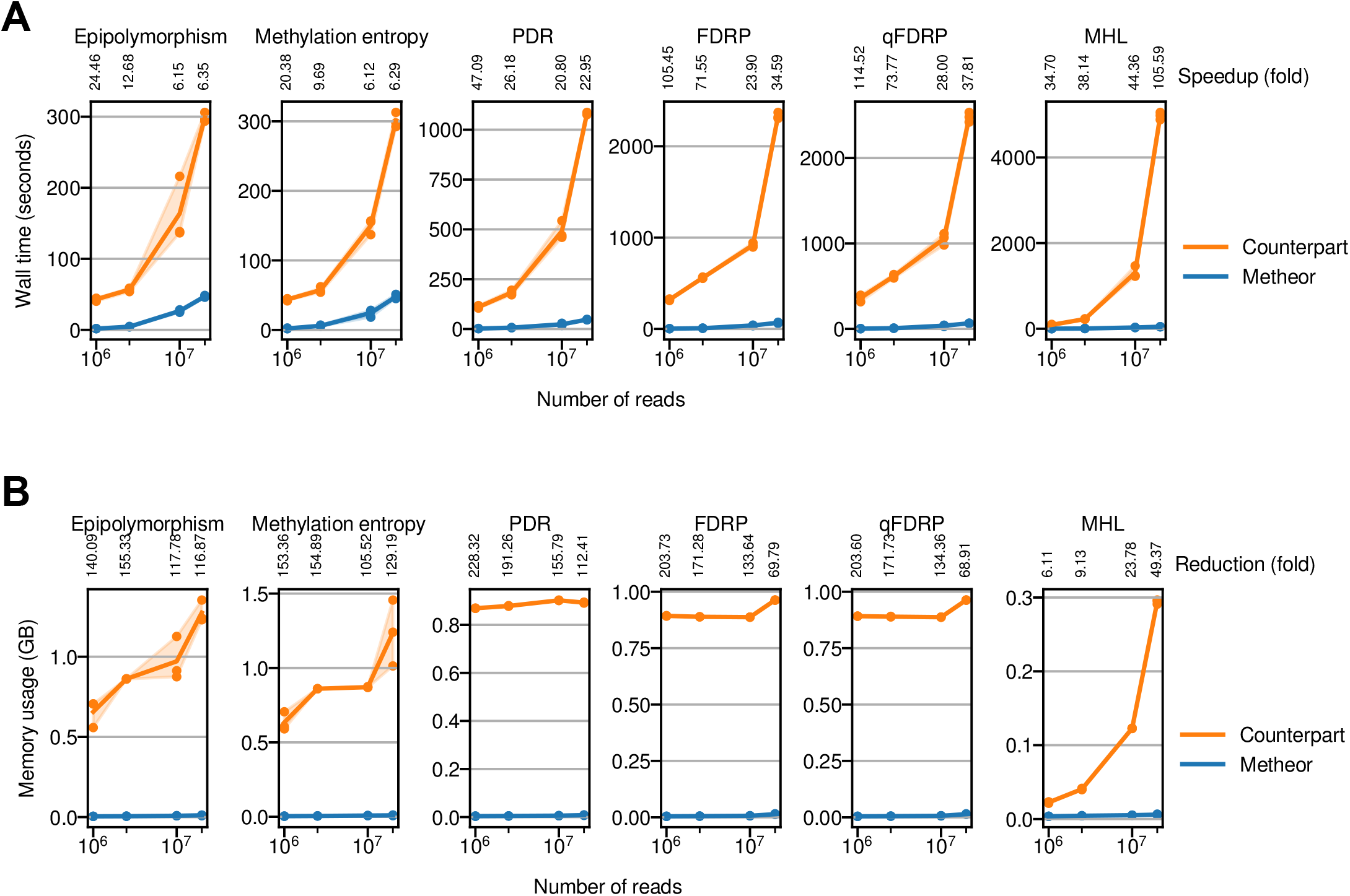
(A) Benchmarking the running time of Metheor using simulated pseudo-WGBS dataset. Values below the name of each of the measures denote the amount of speedup (in fold) in Metheor compared to its benchmark counterpart. (B) Benchmarking the memory usage of Metheor using simulated pseudo-WGBS dataset. Values below the name of each of the measures denote the amount of memory usage reduction (in fold) in Metheor compared to its benchmark counterpart. All the benchmarking experiments were repeated for three times, except for MHL. Lines denote the average wall time and shades represent the 95% confidence interval. Wall time and memory footprint for MHL computation was measured for only once.

**Supplementary Fig. 4.**
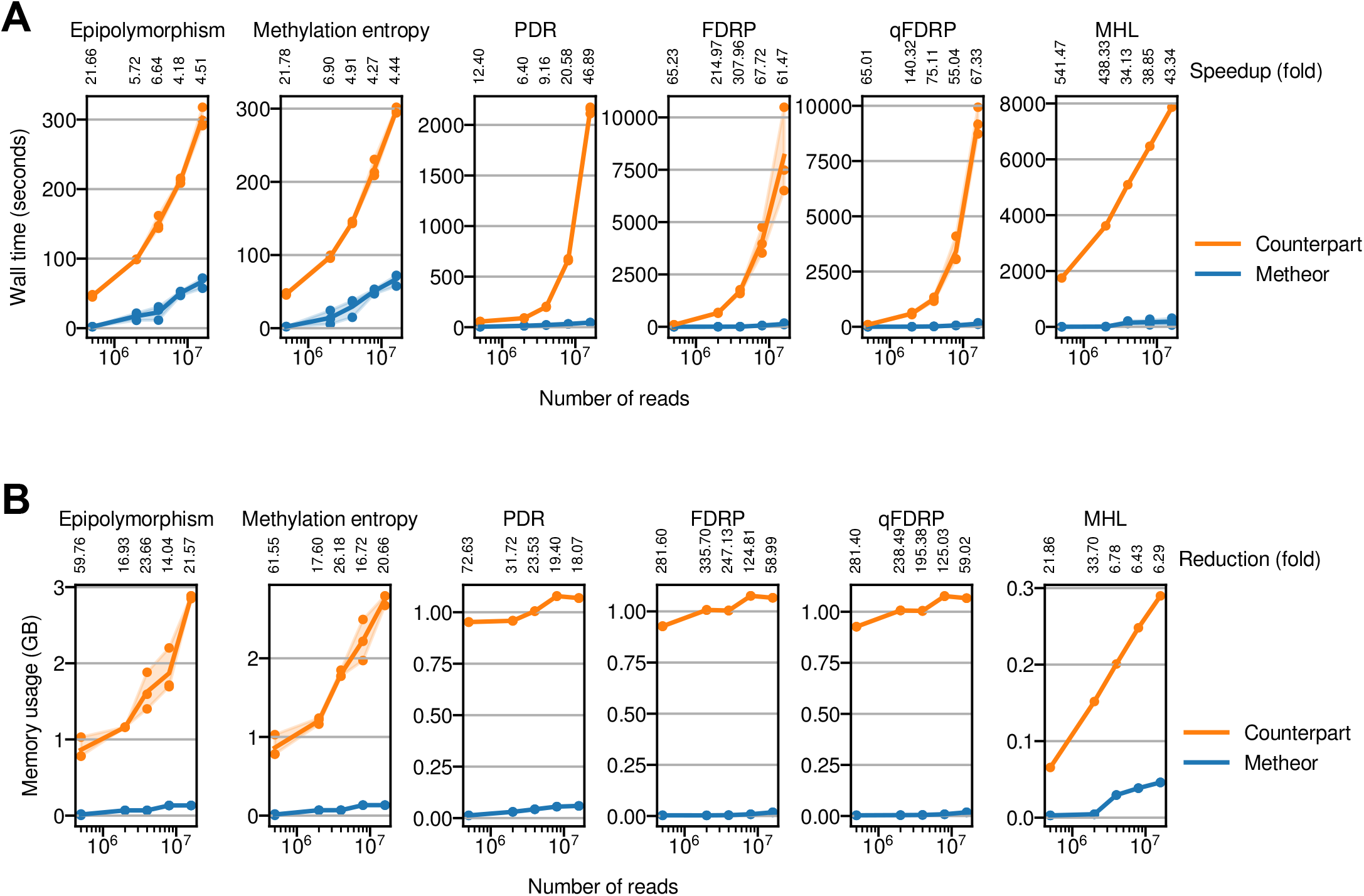
(A) Benchmarking the running time of Metheor using Ewing sarcoma RRBS dataset. Values below the name of each of the measures denote the amount of speedup (in fold) in Metheor compared to its benchmark counterpart. (B) Benchmarking the memory usage of Metheor using Ewing sarcoma RRBS dataset. Values below the name of each of the measures denote the amount of memory usage reduction (in fold) in Metheor compared to its benchmark counterpart. All the benchmarking experiments were repeated for three times, except for MHL. Lines denote the average wall time and shades represent the 95% confidence interval. The wall time for MHL computation was measured for only once.

This merged FASTQ file was processed with Bismark, and the resulting BAM file was used as simulated pseudo-WGBS alignment.

The simulation was done with varying *N* to generate FASTQ files with 1M, 2.5M, 10M and 20M sequencing reads.

#### 1.10 Ewing sarcoma RRBS read subsampling

Finally, we downloaded a real-world Ewing sarcoma RRBS data under SRA run accession SRR5222549 as a benchmark data. The library was constructed from a ewing sarcoma tumor sample and consisted of 18.9M single-end bisulfite reads. To measure the performance of Metheor and the other tools for various sizes of sequencing data, we resampled the sequencing reads to generate differently sized raw sequencing data. As a result, five different raw sequencing data (in FASTQ format) with (1) 500K, (2) 2M, (3) 4M, (4) 8M and (5) 16M reads were generated. Sampled sequencing reads were subsequently adapter-trimmed with trim-galore!, aligned to hg38 reference genome with Bismark, and the resulting alignment files were used for benchmark.

#### 1.11 Benchmarking against WSHPackage using multiple threads

Finally, we compared the running time of Metheor to that of WSHPackage using multiple threads. We particularly focused on the three measures, PDR, FDRP and qFDRP, for which WSHPackage allows the computation to be multithreaded. The running time was measured for 20M simulated RRBS reads, with three replicates using each of 4, 8, 16 and 32 threads for WSHPackage. We observed that Metheor, even with a single thread, was significantly faster than 32-threaded WSHPackage (Supplementary Table 1).

**Supplementary Table 1.** Performance comparison with WSHPackage using multiple threads for 20M RRBS-simulated reads. Values denote the wall time in minutes to compute each measure. Fold speedups in corresponding run of Metheor are shown in parentheses.

| Tool | # threads | PDR (fold) | FDRP (fold) | qFDRP (fold) |
| --- | --- | --- | --- | --- |
| **Metheor** | **1** | **0.87 (1)** | **2.96 (1)** | **3.72 (1)** |
| WSHPackage | 4 | 34.03 (38.92) | 1813.86 (612.65) | 1832.19 (492.70) |
|  | 8 | 19.57 (22.38) | 1072.79 (362.35) | 1097.32 (295.09) |
|  | 16 | 11.21 (12.82) | 652.74 (220.47) | 671.59 (180.60) |
|  | 32 | 11.94 (13.66) | 520.58 (175.83) | 527.40 (141.83) |

#### 1.12 Demonstration of the validity of results

To verify that Metheor produces methylation heterogeneity levels accurately, we compared CpG-wise and CpG quartet-wise methylation heterogeneity levels produced by Metheor to those produced by reference implementations. For experiment, we used the results from simulated RRBS data with 20M reads. Note that PDR, MHL, FDRP and qFDRP levels were computed for each individual CpG, and PM and ME were computed for each CpG quartet. As a result, we observed exactly identical results for PDR, MHL, PM and ME levels (Supplementary Figure 5). For FDRP and qFDRP, values from Metheor and WSHPackage were not exactly the same (Supplementary Figure 5) because these measures depend on random sampling of sequencing reads and it is not possible to reproduce the random process. Nevertheless, they showed extremely high and significant correlation with each other, supporting that Metheor successfully implements procedures to compute FDRP and qFDRP in desired ways.

**Supplementary Fig. 5.**
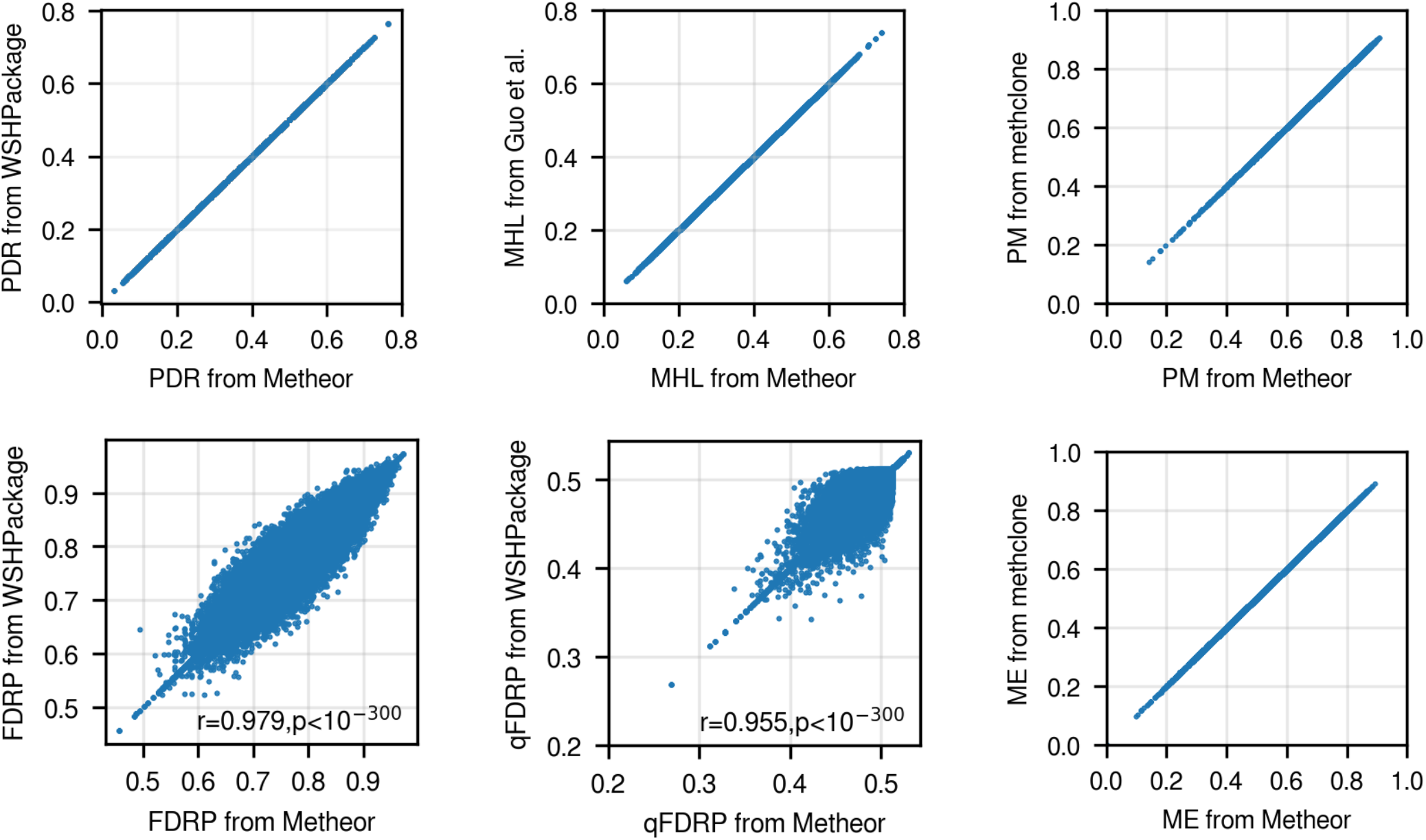
Validity of results. CpG-wise (PDR, MHL, FDRP and qFDRP) and CpG quartet-wise (PM and ME) methylation heterogeneity levels were compared between Metheor and the corresponding reference implementations. Pearson’s correlation coefficient and p-values are shown for FDRP and qFDRP.

